# Antagonistic activity of *Bacillus amyloliquefaciens* subsp. *amyloliquefaciens* against multidrug resistant *Serratia rubidaea*

**DOI:** 10.1101/818054

**Authors:** Sadia Afrin, Mohammad Nazrul Islam Bhuiyan

**Affiliations:** Industrial Microbiology Laboratory, Institute of Food Science and Technology (IFST), Bangladesh Council of Scientific and Industrial Research (BCSIR), Dr. Qudrat-I-Khuda Road, Dhaka-1205, Bangladesh

**Keywords:** Antagonistic activity, multidrug resistance, zone of inhibition, *Bacillus amyloliquefaciens*, *Serratia rubidaea*

## Abstract

*Serratia rubidaea* a member of the Enterobacteriaceae family, is a Gram-negative opportunistic pathogen, known to survive harsh environmental conditions and responsible for hospital associated infections. Specifically, *S. rubidaea* can withstand desiccation and survive on hospital surfaces and equipments as well as have acquired antimicrobial resistance determinants for different commercial antibiotics. The expansion of this multidrug resistant pattern suggests that the treatment of *S. rubidaea* infections will become increasingly difficult in near future. Although some measures were taken to control this species, an inhibition mechanism is remaining unknown. To design effective means to control the dissemination of *S. rubidaea*, an in-depth analysis is required. In the present study, one possible candidate was isolated from the soil of Sundarban Mangroove Forest (Bangladesh) that has important physiological effects to inhibit this pathogenic bacterium. The bacterial isolate was initially identified as *Bacillus amyloliquefaciens* subsp. *amyloliquefaciens* using BIOLOG™ identification system and confirmed to be *B. amyloliquefaciens* strain through 16_S_ rDNA sequence analysis. The growth and antagonistic activity of this potential strain was shown to be stable under wide range of pH, temperature and salinity (NaCl). Moreover, the novel *B. amyloliquefaciens* isolate can also inhibit *Staphylococcus aureus, Escherichia coli, Pseudomonas aeruginosa* and other pathogenic bacteria. These results suggest that *B. amyloliquefaciens* might have potential antimicrobial properties and further research is required for future use of this bacterium as biological controls of *S. rubidaea* or development as new drugs for pathogenic bacteria.

## 1 Introduction

The genus *Serratia rubidaea* is gram-negative speculator pathogens, known to survive harsh environmental surroundings and is a leading cause of hospital acquired infections. This species was first described as *Bacterium rubidaea* in 1940 and presently recognized as *S. rubidaea* [1]. *Serratia*s are widely distributed saprophytic bacterium and has been found in soil, air, water as well as plants. This bacterium is usually responsible for human opportunistic infections [2]. It can cause infections in severely ill patients receiving broad-spectrum antimicrobials or those that have undergone surgery or other invasive procedures including sepsis, bacteremia, urinary tract infections (UTIs), wound and pulmonary infections [3, 4]. Treatment of *S. rubidaea* infection is often complicated by multidrug-resistant phenotypes. This species has essential resistance to several antimicrobial groups, including some classes of beta-lactams and tetracyclines [1, 2, 5]. The expansion of this multidrug resistant pattern suggests that the treatment of *S. rubidaea* infections will become increasingly difficult in the future. At present, spreading of antibiotic resistant bacteria is becoming a major public health problem in all over the world [6, 7]. So here is an urgent need to search new compounds or the sources of new compounds to control *S. rubidaea*. Although some measures were taken to clinically control this species, except an inhibition mechanism is remaining unknown.

Biological control by using antagonistic bacteria is widely expected to become an alternative method for the prevention and control of bacterial diseases. Microbial antagonism is a biological process in which certain microorganisms suppresses the growth of other microorganisms through competition for nutrients or secreting the inhibitory substances [8, 9]. In an antagonistic relationship, several microorganisms can indirectly inhibit other microorganisms or reduce their growth by changing pH, osmotic pressure and surface tension. In another way, a number of microorganisms can directly restrain other microorganisms through producing bacteriocin, antibiotic, toxic components and/or antimicrobial substances [10, 11]. Numerous studies have suggested that these substances are typically antimicrobial proteins or peptides that inhibit the growth of sensitive bacterium or execute them by interfering with the synthesis of cell wall or by forming pores in the cell membrane [12-14]. In a study, J.G. Burgess et al reported about 35% of surface-associated bacteria of Scottish coastal seaweed and invertebrate species that produce antimicrobial compounds [15]. Furthermore, Hans-Peter Grossart *et al*., also reported about several isolates from the Southern Californian Bight that showed antagonistic activity towards other bacteria [16]. In recent years, many studies have been emerged the antimicrobial properties of the genus *Bacillus*. The Gram positive, aerobic, rod-shaped endospore forming bacteria of the genus *Bacillus* are most notably produced antimicrobial substances as secondary metabolites [17]. Shelar *et al*., determined that antimicrobial substances produced by *Bacillus atrophaeus* JS-2 showed antagonistic activity against both gram-positive and gram-negative microorganisms [18]. The Gram positive bacterium *Bacillus subtilis* is a vital producer of industrial enzymes and can also able to demonstrate antagonism behavior to other bacteria. Recently, Das *et*.*al*., isolates one novel *Bacillus subtilis* AN11 strain from Bhitarkanika mangrove forest, Odisha, India that have strong antagonistic activity against major fish bacterial pathogens [8]. These observations suggest the involvement of *Bacillus* genus in responses to pathogen resistance processes.

*Bacillus amyloliquefaciens* is a non-pathogenic soil bacteria closely related to the species *B. subtilis* [19]. This bacterium is gram positive rods along with peritrichous flagella consent to motility. The cells often appear as long chains unlike many other *Bacillus* species that form as single cells. Similar to other *Bacillus* species, *B. amyloliquefaciens* forms endospores allowing continued existence for a long time period. Endospores appear centrally in the cells which do not have a swollen appearance [20, 21]. This species have impressive capacity to synthesize non ribosomal secondary metabolites with antimicrobial activity [22, 23]. On the basis of phylogenic analysis and physiological characteristics, recently *B. amyloliquefaciens* divided in to two sub species: the plant associated *B. amyloliquefaciens* subsp. *plantarum* and non plant associated *B. amyloliquefaciens* subsp. *amyloliquefaciens* [24]. In the present study, one novel *B. amyloliquefaciens* subsp. *amyloliquefaciens* species was isolated from the soil of Sundarban Mangrove Forest, Bangladesh. The objectives of this study were to evaluate the antagonistic activity of *B. amyloliquefaciens* subsp. *amyloliquefaciens* against *S. rubidaea*. The inhibition profiles of *B. amyloliquefaciens* subsp. *amyloliquefaciens* were conducted in response to different temperature, pH and salinity (NaCl). Furthermore, antagonistic spectrum of *B. amyloliquefaciens* was analyzed using various bacterial pathogens. To the best of our knowledge, this is the first report on a promising antagonistic *B. amyloliquefaciens* subsp. *amyloliquefaciens* strain from Sundarban Mangrove Forest soil in Bangladesh against *S. rubidaea*. The obtained results give new insight in the development of bio-controls agent to prevent and control *S. rubidaea* infections.

## 2 Materials and Methods

### 2.1 Collection of soil samples

Sundarban Mangrove Forest of Bangladesh was selected for the collection of soil samples. Soil samples were collected in March, 2018 from different locations. Collected soil samples were kept in sterilized plastic bags and immediately brought to the laboratory and was preserved in refrigerator before and after analyses. The pH of the soil samples were determined by using a digital pH meter (Jenway 3310 pH meter, U.K.) making soil paste at 1 : 2 (Soil : Water) ratio.

### 2.2 Screening of antagonistic bacteria

Once delivered to the laboratory, samples were taken to the procedure for isolation. Pour plate technique was used to isolate the desired organisms. Samples (1 gm) were used directly and also diluted to 10^−1^, 10^−2^, 10^−3^, 10^−4^ and 10^−5^ using sterile distilled water. Different samples and dilutions were plated into Muller Hinton Agar (MHA) medium (HiMedia) and Tryptic Soya Agar (TSA) medium (HiMedia). The plates were incubated at 37°C for 24-48 h. Based on inhibition zone discrete bacterial colonies were selected for isolation and were purified by repeated streaking method. Isolated colonies were preserved in a refrigerator at 4°C. The identification of these bacterial strains was carried out on the basis of morphological characteristics, Gram staining characteristics, BIOLOG™ identification system and PCR identification system.

### 2.3 Morphological Characterization

Antagonistic isolates were purified by continuous subculture on Nutrient agar (NA) medium (HiMedia). Colony morphologies (color, shape and size) and microscopic observations were examined after 24 h incubation. The morphological characteristics of the isolates were checked by Gram staining procedure [25].

### 2.4 Identification by BIOLOG™ System

For the species-level identification of antagonistic isolates, BIOLOG™ identification system was applied (BIOLOG™, USA) base on the utilization of 71 carbon sources and 23 chemical sensitivity assays in GEN III microplate test panels. The isolates were cultured on biolog universal growth (BUG) agar medium. All microplates and inoculating fluid were pre-warmed at 37°C for 30 mins. After 18 h incubation the inoculum of antagonistic bacterial isolates were added to the inoculating fluid-A for obtaining the desired turbidity which is (90-98) % T because the target cell density must be in the range of (90-98) % T for protocol-A of GEN III Microbial ID 20 assay techniques on GEN III microplate. The cell suspension was then poured into the multichannel pipette reservoir and the tips of multi-Channel. Repeating Pipettor was filled by drawing up the cell suspension from the reservoir and all 96 wells were inoculated with accurately 100μl of bacterial suspension. The microplate was then covered with its lid and incubates at 37°C for 18 to 24 h. After incubation, the microplate was placed into the Micro Station Reader for analysis to obtain ID result and the result was given through comparing with database using the software program MicroLog 4.20.05 (BIOLOG™, USA) [26]. The scope of the 96 assay reactions, coupled with sophisticated interpretation software, delivers high level of accuracy that is comparable to molecular methods.

### 2.5 DNA Extraction and PCR amplification of antagonistic bacterial

Heat-thaw method was applied for the DNA extraction [27]. Isolates were grown in Nutrient broth (NB) (Difco™) at 37°C for 24 h. 1ml broth culture was taken into eppendrof and centrifuged for 5 mins at 10000 rpm. Following, pellet was collected then 100µl RNase free water (Thermo Fisher Scientific) was added and uniformly mixed by vortexing. At 100°C in water bath the eppendrof tubes were boiled for 10 mins and immediately after boiling the tubes were placed into ice for another 10 mins then the tubes were centrifuged for 5 mins at 10000 rpm. The supernatant was collected and store at −20°C which contains bacterial DNA.

For PCR amplification, the reactions were carried out in 30µl mixture composed of 22.5µl PCR MasterMix (Invitrogen), 1.2 µl of each forward and reverse primer (primers F27/ R1492), DNA template 2µl and 3.1 µl RNase free water (Thermo Fisher Scientific) 35 cycles using a thermal cycler (BioRad) and amplification conditions were 94°C for 1 min, 60°C (for primers F27/ R1492) for 1 min, and extension at 72°C for 1 min followed by final extension was conducted at 72°C for 5 min using a PCR minicycler (Eppendorf Ltd., Germany). After amplification, PCR products were investigated with 1.0% agarose (Invitrogen) gel in 1X TAE buffer (50X TAE buffer, Thermo Fisher Scientific) by electrophoresis (Compact XS/S, Biometra) and DNA bands were visualized with ethidium bromide (Thermo Fisher Scientific) under ultraviolet (UV) transilluminator (Biometra).

### 2.6 Antibiotic resistance profiles of *S. rubidaea*

The selected strain was investigated for their antibiotic resistance profiles as recommended by the Clinical and Laboratory Standards Institute (CLSI; Wayne, PA, USA). The strain was grown overnight (24 h) into NB medium at 37°C. Petri dishes containing 20 ml of MHA medium allowed to solidify at room temperature and swabbed with inoculated broth. In this study, 18 (eighteen) different types of antibiotic discs such as Penicillin G (10 µg), Ampicillin (10 µg), Vancomycin (30 µg), Erythromycin (15 µg), Imipenem (10 µg), Amikacin (30 µg), Doxycycline (30 µg), Gentamicin (10 µg), Nalidixic acid (30 µg), Nitrofurantoin (300 µg), Neomycin (10 µg), Tetracycline (30 μg), Cephotaxime (30 µg), Kanamycin (30 µg), Ceftazidime (30 µg), Ciprofloxacin (5 µg), Metronidazole (5 µg) and Tobramycin (10 µg) were used. The agar plates were incubated at 37°C for 24 h. Diameter (in mm) of the inhibition zone was measured using an antibiotic zone scale. Inhibition zones diameters of antibiotics were compared to those defined by Charteris *et al*., 1998 [28] and results were expressed in terms of resistance, moderate susceptibility or susceptibility by comparing with the interpretative zone diameters provided in the performance standards for antimicrobial disc susceptibility tests.

### 2.7 Antagonism assay

For the determination of growth and antagonistic activity of these isolates, three major selection criteria were chosen such as different temperatures, a range of pH and different concentrations of salinity (NaCl) [29]. For this purpose, active cultures (incubated for 16-18 h) were used.

#### 2.7.1 Thermal stability assay

NB medium was prepared and transferred to tubes in 5 ml. Then 50μl cultures inoculated to that tubes and incubated for 24 h at 15°C, 20°C, 25°C, 30°C, 35°C, 37°C, 40°C, 45°C, 50°C and 55°C. After these incubation periods, cells growth at different temperatures was observed and recorded. Growth was monitored at OD 600 nm (Thermo Multiskan EX). At the same time antagonistic activity of the isolates were also observed by the agar well diffusion method.

#### 2.7.2 pH stability assay

The influence of pH on the growth and antagonistic was measured with varying range of pH from 4.5, 5.0, 5.5, 6.0, 6.5, 7.0, 7.0, 7.5 and 8.0. The low pH value of the NB was adjusted to 1M HCl. The inoculated broths were then incubated for 24 h at 37°C. Antagonistic activities of the isolates were also observed within the same pH range.

#### 2.7.3 Salinity (NaCl) stability assay

The isolates were tested for the tolerance against different NaCl concentrations. The growth rate of bacterial cultures in NB containing different levels (1%, 2%, 3%, 4%, 5%, 6% and 7%) of NaCl was determined. In addition, antagonistic behaviors of the isolates were observed under the identical NaCl concentration.

### 2.8 Minimum inhibitory concentration (MIC) and Minimum bactericidal concentration (MBC) of *B. amyloliquefaciens*

The supernatant of *B. amyloliquefaciens* isolats was filtered through 0.2 μm membranes and used to determine the MIC and MBC. In brief, the supernatant were diluted in 96-well plates by two fold serial dilutions using MHB. Then *S. rubidaea* of approximately 10^5^-10^6^ CFU/ml was added into each well, mixed gently and then incubated at 37°C for 24 h. The last concentration that provided clear solution when compared to the growth control was recorded as the MIC. The MBC was evaluated by pipette each dilution from the clear wells, diluting with PBS pH 7.2 and then 10 μl of each dilution was dropped onto NA medium for colony counts. The interpretation was that the MBC must have decreased ≥3 log^10^ CFU/ml or 99.9% of the bacterial cell count when compared to growth control.

### 2.9 Bio-control of *S. rubidaea* using *B. amyloliquefaciens*

*S. rubidaea* at 10^5^-10^6^ CFU/ml (approximately) were co-cultured with 10^5^-10^6^ CFU/ml (approximately) of *B. amyloliquefaciens* isolate in NA medium in Erlenmeyer flasks (250 ml) and incubated for 24 h at 37°C with 200 rpm shaking. The experiment was done in triplicate. *S. rubidaea* cultured in the same condition was used as a control. The viability of *S. rubidaea* was measured after 24, 48, 72 and 96 h using the plate count method on NA medium, respectively.

### 2.10 Antimicrobial activity of *B. amyloliquefaciens* against others pathogenic bacteria

The selected strain was investigated for antimicrobial activity against a variety of pathogenic microorganisms. Agar well diffusion method was modified and used to detect antimicrobial activities [30, 31]. These assays were performed in triplicate. The plates were poured with 20 ml MHA media. Nine different pathogens belonging to both gram-positive and gram-negative groups such as *Bacillus cereus* ATCC 10876, *Bacillus subtilis* ATCC 11774, *Staphylococcus aureus* ATCC 9144, *Escherichia coli* ATCC 11303, *Salmonella typhi* ATCC 13311, *Shigella flexneri* ATCC 12022, *Enterobacter faecalis* ATCC 29212, *Vibrio parahemolyticus* ATCC 17802, *Pseudomonas aeruginosa* ATCC 27853. The pathogenic strains were grown in NB medium for 24 h and spread on the surface in MHA plate. Three wells of each plate were made by using a sterile borer and 10 µl of the supernatant of the *B. amyloliquefaciens* were placed into three well. The plates were preinoculated at room temperature for the diffusion and incubated overnight at 37°C for 24 h. The plates were examined for zones of inhibition. At the end of the incubation, inhibition zone diameters were measured.

### 2.11 Partial analysis of bacterial metabolites

Protein was precipitated according to the Acetone precipitation method [32]. Broth media with *B. amyloliquefaciens* cells were centrifuge at 10,000 rpm for 10 min at 4°C and supernatant were collected. Ice cold (−20°C) acetone was added three times (150µl) the sample volume (50µl) to the ependrof tube. After mixing well and incubating at −20°C for overnight, centrifugation was done at 13,000-15,000 rpm for 30 min at 4°C temperature. Supernatant was disposed and the acetone was allowed to evaporate from the uncapped tube at room temperature for 30 minutes. 200µl of SDS buffer was added to the protein pellet and to determine the protein profiles extracted though Acetone precipitation method, SDS-PAGE analysis (Bio-Rad, USA) was done with 5% stacking and 10% separating gel at 20 mA followed by comassie blue staining dyes. After the run had been completed, the gel was stain and destains in 0.1% Coomassie Brilliant blue and 10% Acetic acid for approximately 2 h at 100 rpm.

### 2.12 Statistical Analysis

All experiments were carried out in triplicate. Data were presented as the mean ± standard deviation (SD) for the indicated number of independently performed experiments. *P <* 0.05 was considered statistically significant using one-way analysis of variance (ANOVAs).

### 3.0 Results and Discussion

#### 3.1 Isolation, purification and morphological determination

Soil samples were properly diluted with sterile water and plated into MHA medium thus incubated at 37°C. After 24 h incubation bacterial growth with zone-inhibition assay was observed. According to Lategan *et al*. and Haipeng Cao *et al*., zones of inhibition >12 mm against other organism were considered as susceptibility to this isolate [22, 33]. In the present study, only one isolate was found to exhibit strong antagonistic activity to the other organism, displaying 29 mm inhibition zones. (Figure 01). Thus, the isolates (both resistant and sensetive) were chosen for further study. The selected isolates were observed by optical microscope to determine their morphology. The microscopic study of the isolates showed the resistant strain to be Gram-positive rods, where the sensitive bacteria designated itself as Gram-negative coccoid rod-shaped less than 1 μm in diameter. Further microscopic observation demonstrated that the resistant strain sporulated aerobically without swelling of cell, indicated that tested strains belong to the genus *Bacillus* and presumptively resistance to high temperatures. On the other hand the sensitive strain does not produce spore.

**Figure 01:**
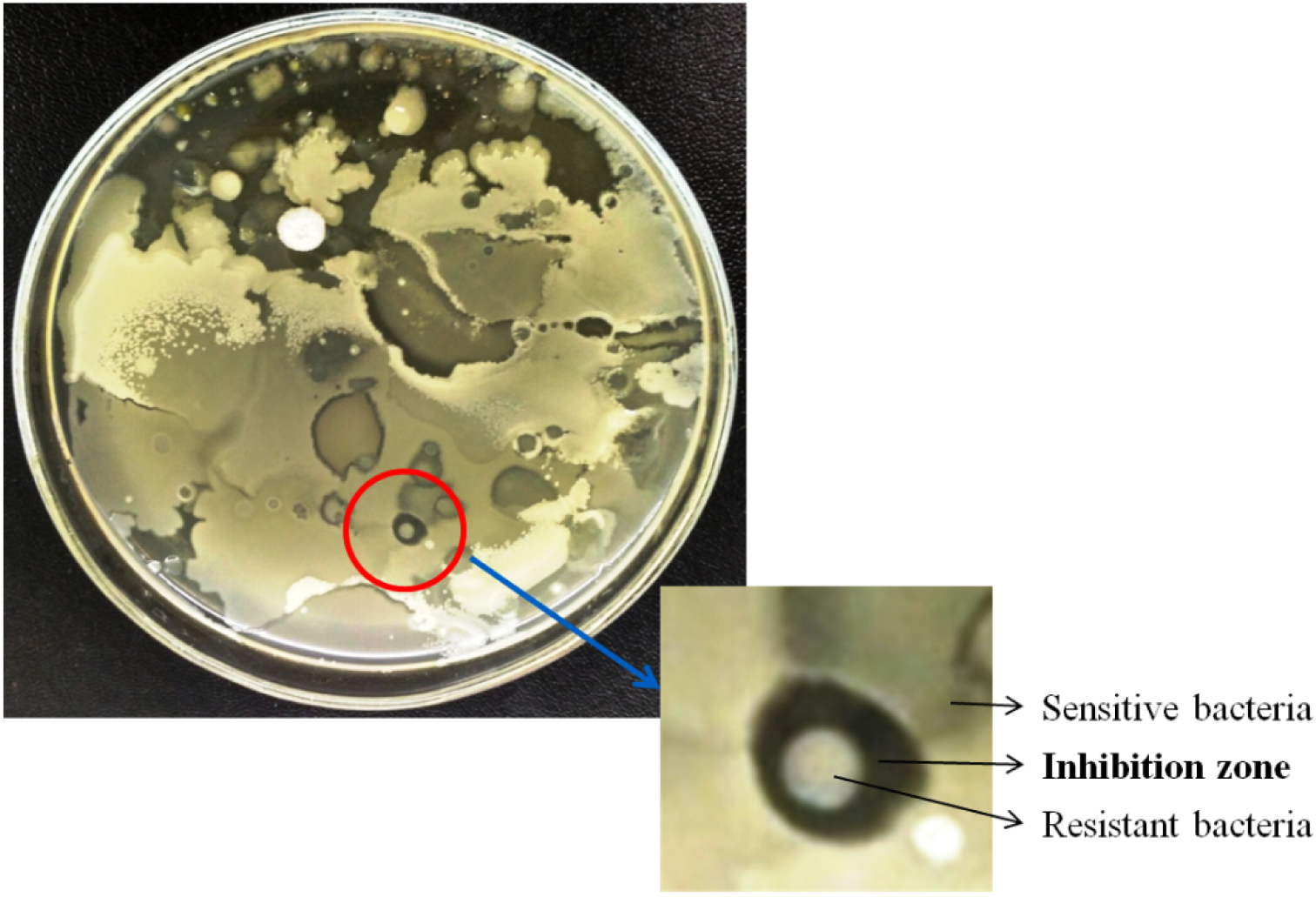
Bacterial colonies from soil suspension.

### 3.2 Identification of isolates using BIOLOG™ system

The strains were correctly identified with BIOLOG™ system up to the species level. The BIOLOG™ system aims to provide a rapid, convenient approach to bacteria identification with a database of 3000 species. The result indicates that the resistant strain identified as *Bacillus amyloliquefaciens* ss *amyloliquefaciens* and the sensetive strain was identified as *Serratia rubidaea*. For further confirmation these isolates examined for three replications. Table 01 shows the result of BIOLOG™ identification system.

**Table 1:** Microorganism identification with BIOLOG™

| Strain | ID<br>(identification) | PROB<br>(probability) | SIM<br>(similarity<br>index) | DIST<br>(distance) |
| --- | --- | --- | --- | --- |
| Resistant<br>strain | *Bacillus amyloliquefaciens*<br>ss *amyloliquefaciens* | 0.850 | 0.561 | 4.854 |
| Sensitive<br>strain | *Serratia rubidaea* | 0.674 | 0.674 | 4.683 |

### 3.3 Molecular Identification

For additional confirmation, these isolates were further identified using RT-PCR (Reverse transcription) method. PCR amplification against the conserved regions in 16s rDNA genes of resistant strain gave the 1400 base pair (bp) products (Fig. 2A) where 1450 bp amplicon was observed for the sensitive strain (Fig. 2B). In the previous study, A.M. Hanafy *et al*., 2016 reported about *B. amyloliquefaciens* that was around 1400 bp long and this data is very much consistent to our present result [34]. Similarly, the data of RA Bonnin et al 2015 support the sensitive isolate identification as *S. rubidaea* [2].

**Figure 02:**
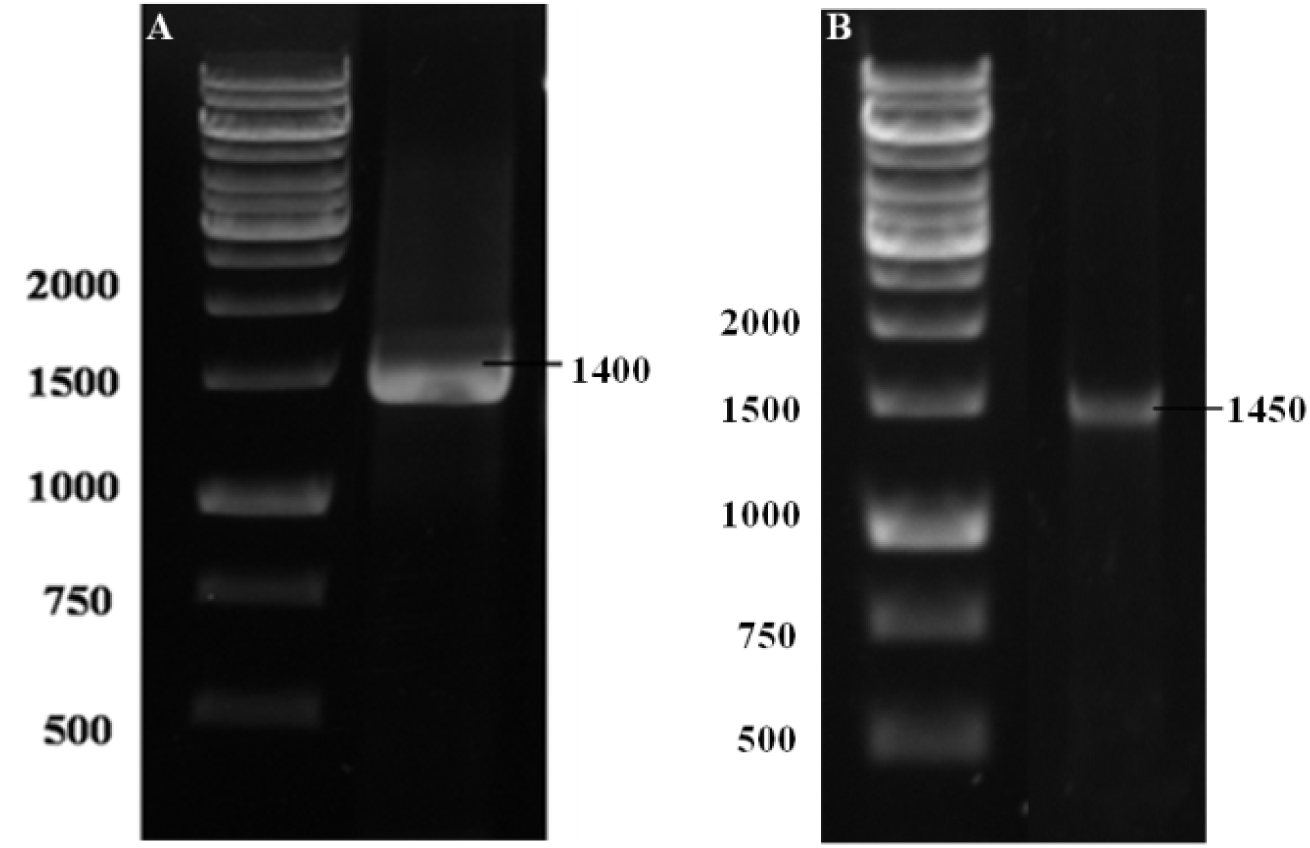
Agarose gel electrophoresis. A. 1400 bp size band of resistant strain, B. 1450 bp size band of sensitive strain. DNA ladder: 1Kb.

### 3.4 Multidrug resistance pattern of *S. rubidaea*

Increasing evidences suggest that the speculator bacteria *S. rubidaea* exhibit a resistant profile toward a number of commercial antibiotics that leads major health problem and seriously threatens to control this bacteria [35, 36]. To determine the resistance patterns of *S. rubidaea*, culture and sensitivity test was performed. In the present experiment, eighteen (18) commercial antibiotics were used to compare the zone of inhibition. According to this test the isolate of *S. rubidaea* was defined as multi drug resistance (MDR) strain as they were resistant to three or more unique antimicrobial classes: including aminoglycosides, quinolones, extended-spectrum cephalosporins, beta-lactam/beta-lactamase inhibitor combination (Fig 03 and Table 02).

**Table 2:**
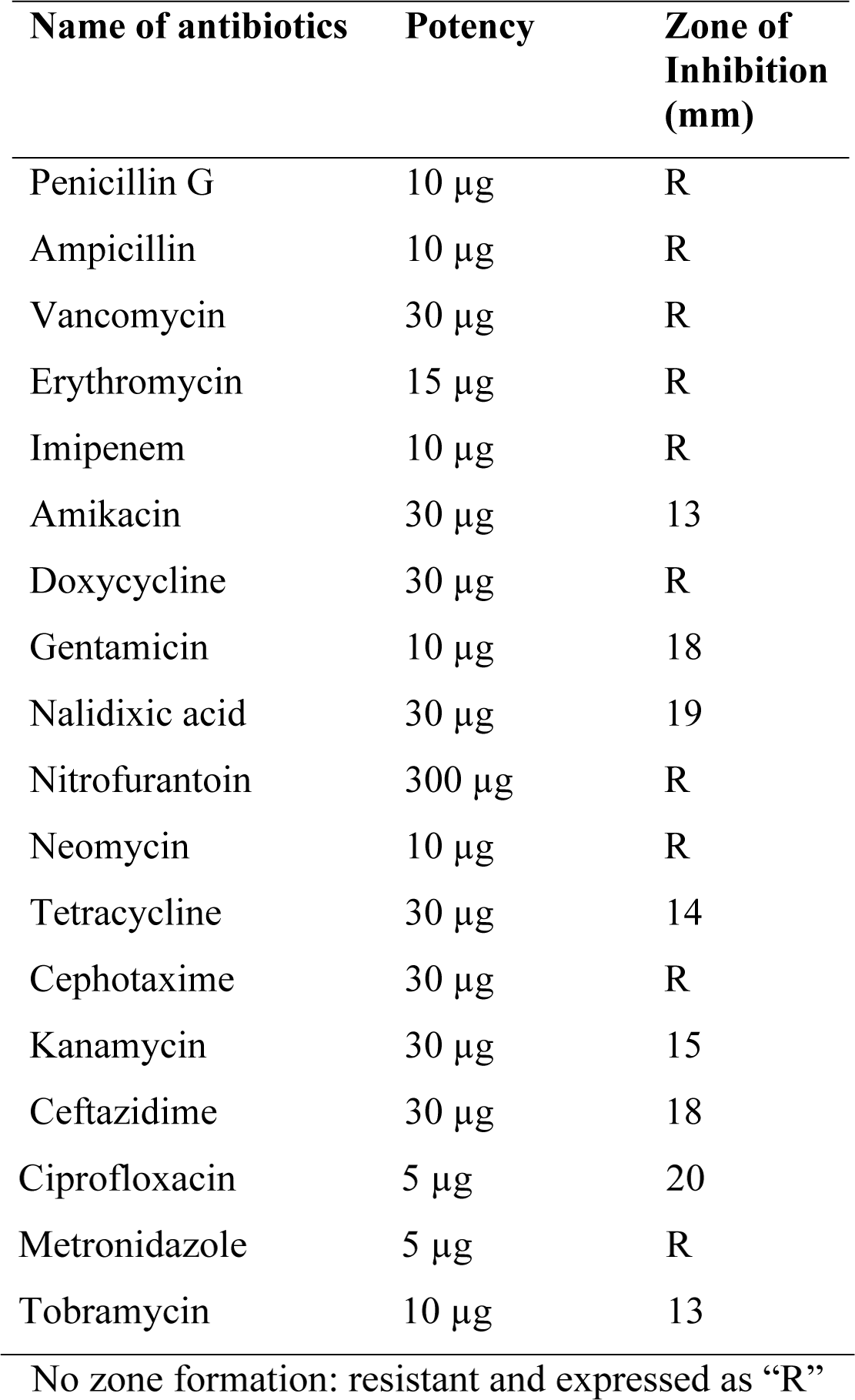
Resistance or susceptibility of *S. rubidaea* to eighteen (18) commercial antibiotics

**Figure 03:**
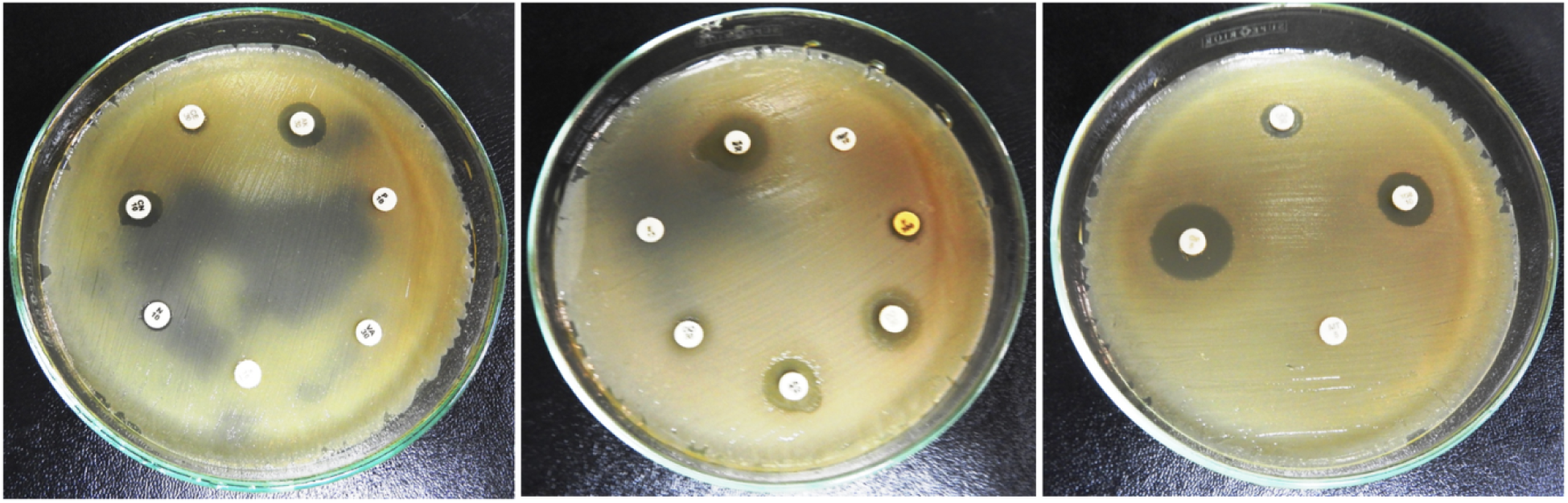
Antibiotic sensitivity test of *S. rubidaea*. Inhibition zone: sensitive, no zone formation: resistant.

### 3.5 Inhibition potentiality tests of *B. amyloliquefaciens* against *S. rubidaea*

Agar well diffusion method is widely used to evaluate the antimicrobial activity [37]. In this present study, well diffusion method with 6 mm diameter was used to evaluate the potentiality of *B. amyloliquefaciens* against *S. rubidaea*. The MHA plate is inoculated by spreading a volume of *S. rubidaea* inoculums over the entire agar surface. Then, a hole is punched aseptically with a sterile cork borer and a volume (20-100 µl) of the *B. amyloliquefaciens* inoculum is introduced into the well. After that, agar plates are incubated at 37°C for 24 h. The antimicrobial agent of *B. amyloliquefaciens* diffuses in the agar medium and inhibits the growth of the *S. rubidaea* (Fig.4A). For further confirmation of the potentiality of *B. amyloliquefaciens*, cross test was also performed (Fig.4B) that indicating *B. amyloliquefaciens* may have some active substances which inhibit the growth of opportunist *S. rubidaea*. As a next step, the whole above mention procedure was done for TSA medium, which gives the identical outcome as MHA medium.

In order to further understand the activities of *B. amyloliquefaciens* against *S. rubidaea*, the mode of action of whole bacterial cell culture and the cell free supernatant (CFS) was recorded separately. For the preparation of CFS, bacterial culture was incubate in broth media and harvested by centrifugation at 8500 rpm for 10 minutes at 4°C. To determine the antimicrobial spectrum, whole bacterial cell (100μl) and CFS (100μl) were placed in 6mm diameter well on both MHA and TSA media. After 24 h of incubation results were observed and recorded in table 03. This experiment was repeated for three times. The result indicated that CFS of *B. amyloliquefaciens* can restrain the growth of *S. rubidaea* where as whole bacterial cell more actively suppress the growth of *S. rubidaea* with in the same time period (Fig. 4C and 4D).

**Table 3:** Inhibition zone (mean±SD) of *B. amyloliquefaciens* against *S. rubidaea* in different media

| Media | Inocula type |  |
| --- | --- | --- |
|  | Bacterial culture | Cell free supernatant |
| Inhibition zone (mm) on MHA | 29.0±0.24 | 19.0±0.27 |
| Inhibition zone (mm) on TSA | 26.5±0.22 | 18.0±0.28 |

**Figure 04:**
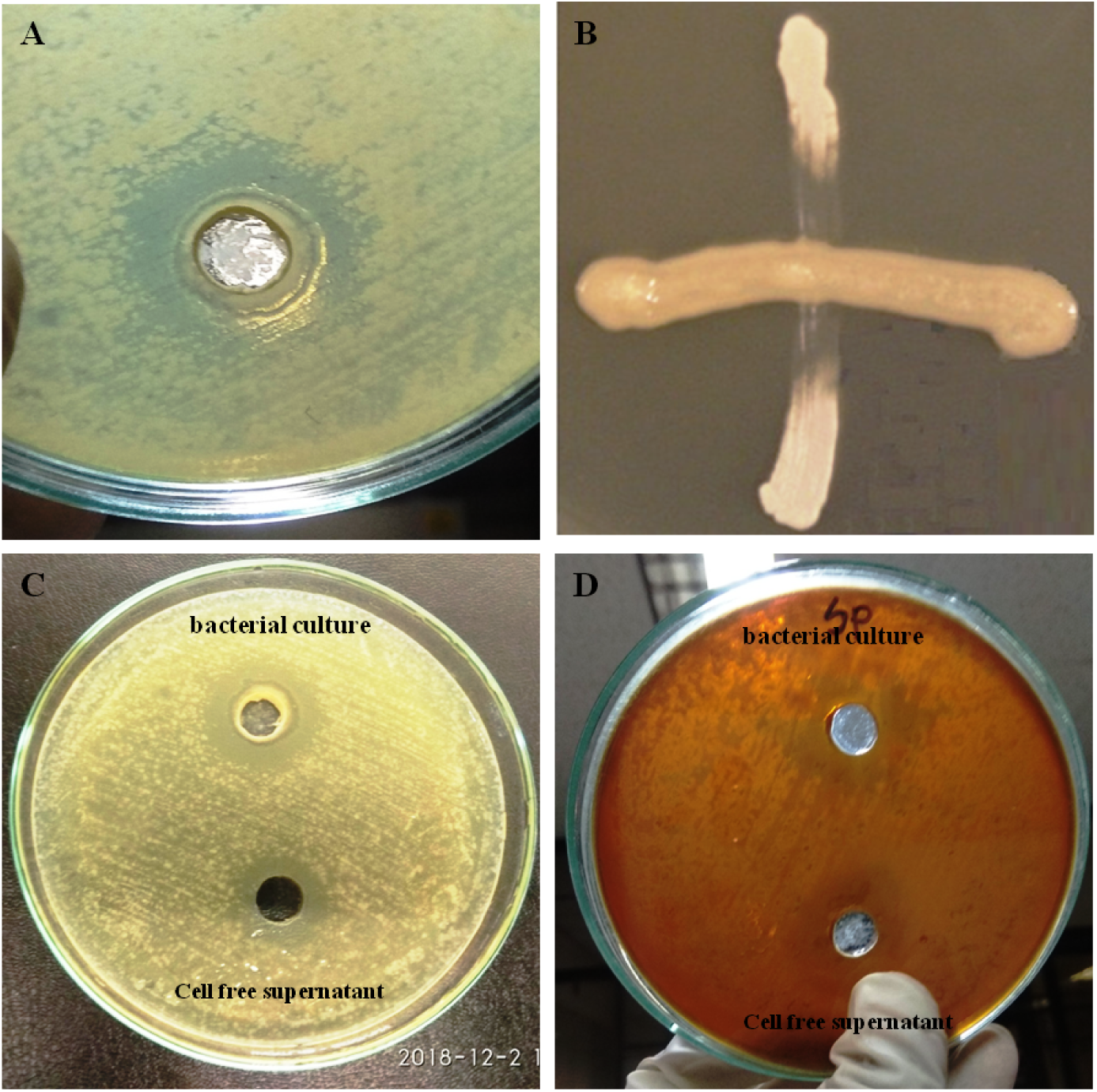
**A**. Inhibition potentiality of *B. amyloliquefaciens* against *S. rubidaea*, B. Cross test of horizontally streaked *B. amyloliquefaciens* bacterium across vertically streaked *S. rubidaea* resulting in inhibited *S. rubidaea* growth, C. Inhibition zone (mm) of whole bacterial cell and cell free supernatant on MHA, D. Inhibition zone (mm) of whole bacterial cell and cell free supernatant on TSA.

### 3.6 Physiological parameters on growth of *B. amyloliquefaciens*

Morphological feature of *B. amyloliquefaciens* indicating that, the bacterial colony was light gray in color, opaque with a rough matted surface and having irregular edge. The earlier studies reveled that most of the antibiotics and/or antibiotic like substances were formed both during spore formation and by actively multiplying cells [38]. In this present study, we found that the bacterium *B. amyloliquefaciens* is very fast growing and has an optimum growth after 16 h that produce spore after 24 h of incubation. Important biochemical tests *viz*. carbon and nitrogen source utilization, geletin liquefaction test, voges proskauer (VP) test and H_2_S tests were carried out (Table 03) for recognizing the nature of *B. amyloliquefaciens*. The result of biochemical tests indicated that, the selected strain can utilize majority of carbon and nitrogen sources. The previous studies specify that *Saccharomyces* yeast exhibit negative result in both indole and H_2_S production [19, 24]. The present study also comply the same phenomenon as previous studies.

Optimization of culture conditions is very important for maximum microbial growth as well as inhibition capacity of microorganisms [39, 40]. Among the physical and chemical parameters, optimum temperature, pH range and salt concentration are the most important as well as the time period is also very significant [41, 42]. To acquire maximum growth of *B. amyloliquefaciens*, optimum growth condition was confirmed by the measurement of optical density (OD) at 600 nm (OD_600_). As showed in Fig. 05, the growth rate of isolate was expressed constitutively at different temperatures with the highest level at 37°C. The organism can able to grow at 50°C but unable to grow below 20°C (Fig. 5A). In the case of pH tolerance test, the growth of the isolate decrease at low pH and found to the maximum growth rate at pH 6.5 as well as the growth declined at pH 8.0 (Fig. 5B).Tolerance to sodium chloride was determined by testing their ability to grow in the presence of different concentrations of NaCl (1.0%, 2.0%, 3.0%, 4.0%, 5.0%, 6.0% and 7.0%). According to the test result, the isolate was grows up to 7.0% NaCl concentration but there was a rapid decrease in growth after 3.0% NaCl concentration (Fig. 5C). Furthermore, to find out the optimum growth at different time periods the representative isolate *B. amyloliquefaciens* was observed at various time phase. The maximum growth occurred during the early stationary phase (24 h). During the extended stationary phase of incubation, the activity of the organism decreased considerably to a complete termination of activity (72 h) (Fig. 5D).

**Figure 05:**
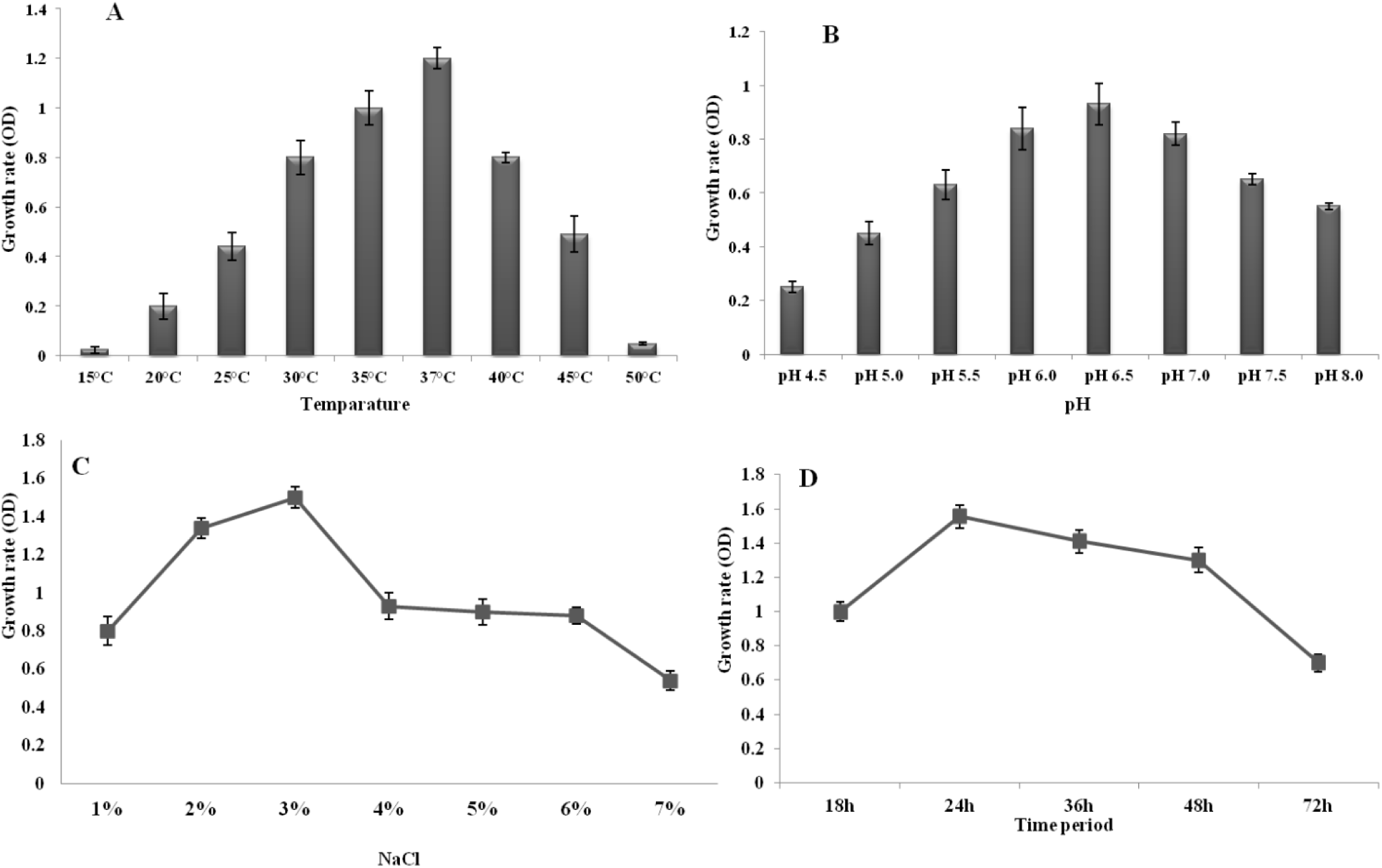
Effect of temperature, pH, NaCl and time period on the growth of *B. amyloliquefaciens*.

### 3.7 Inhibition profiles of *B. amyloliquefaciens* against *S. rubidaea* in response to different temperature, P^H^ and salinity

The use of antagonistic bacteria is widely expected to become an alternative method for the prevention and control of bacterial diseases. *B. amyloliquefaciens* are now recognized as a bacterium with an impressive capacity to synthesize secondary metabolites with antimicrobial activity [43]. Numerous studies have suggest that unusual surroundings such as temperature, P^H^, time period, dose and other matters differentially trigger the inhibition capability of bacteria [16, 44]. Antagonistic activity of *B. amyloliquefaciens* was tested at different temperatures (30°C, 35°C, 40°C, 45°C and 50°C) with in different time (24, 48 and 72h) periods. The temperatures were selected on the basis of optimum growth condition of *B. amyloliquefaciens*. The culture showed prominent growth on selected media at all tested temperature; however, no antagonistic activity against *S. rubidaea* was indicated at 50°C with in 72 h time period (Fig. 6A).

**Figure 06:**
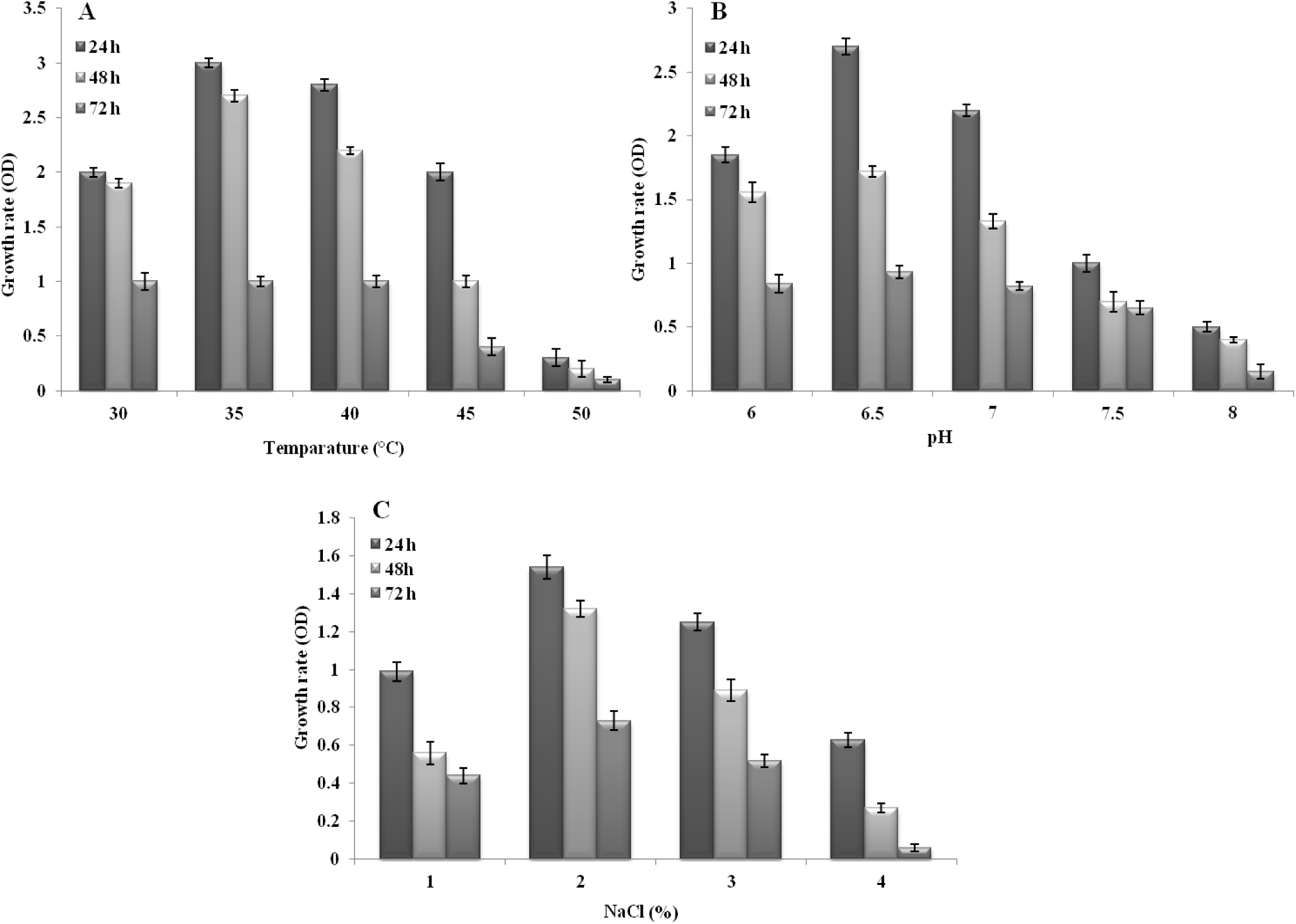
Antagonistic activity of *B. amyloliquefaciens* against *S. rubidaea* within different temperature, pH and salinity.

Among physical parameters, pH of the growth medium plays an important role in inhibition competence of organism. Most of the earlier studies revealed that pH 6.0 to 7.0 is optimum for the growth and inhibition ability of the majority bacterial strains [22, 45]. In this present study, we also found that the pH 6.0 is required for the optimum growth of *B. amyloliquefaciens*. So, the inhibitory effect of *B. amyloliquefaciens* against *S. rubidaea* was investigated at different pH (6.0-8.0). At pH 6.5, more or less fifty percent (50.0%) inhibition of radial growth of *S. rubidaea* was shown by *B. amyloliquefaciens*, which decreased nearly thirty percent (30%) at pH 8.0 (Fig. 6B). So the result reveled that, optimal conditions for antagonistic activity were found to be 37°C and pH 6.5 at 24 h, while the maximum growth was also observed at 37°C and pH 6.5 within the same time period. A shift in temperature and pH below and above this value resulted in considerable reduction in the growth and also the antagonistic activity of *B. amyloliquefaciens* against *S. rubidaea*. However, adequate level of growth and zone of inhibition was still observed at 30°C to 45°C and pH 6.0 to 7.5.

In correspondence to temperature and pH, saltinity (NaCl) enhancement also significantly affect the antagonistic behavior of *B. amyloliquefaciens*. Though the growth of antagonistic isolates was optimally observed up to 7.0% of salt concentration, but the level of inhibition of *B. amyloliquefaciens* against *S. rubidaea* was observed up to 2.0% of salinity level at 24h. Although an increase was observed up to 4.0% of salinity, but no activity was observed against the indicator isolate after 72 h of incubation (Fig. 6C).

### 3.8 Dose dependent inhibition patterns of *B. amyloliquefaciens*

The inhibitory zone produced by *B. amyloliquefaciens* against *S. rubidaea* is shown in Table 05. To evaluate the dose dependent inhibition patterns optimum temperature, pH and salinity were maintained, as well as 24 h incubation period was also determined. The zone sizes were ranged from 8.0 mm to 29.0 mm in various concentrations of bacterial suspension. The highest zone was recorded at 30μl concentration. The MIC value of *B. amyloliquefaciens* recorded as 200 μl/ml against the test bacteria. On the contrary the MBC value noticed at 300 μl/ml against *S. rubidaea* (Table 06).

**Table 4:** Biochemical characteristics of the *B. amyloliquefaciens* strain

| Test | Result | Test | Result |
| --- | --- | --- | --- |
| Glucose | + | Starch hydrolase | + |
| Sucrose | + | Indole production | - |
| Maltose | + | Oxidase | - |
| Mannitol | + | Catalase | + |
| Xylose | + | VP reaction | + |
| Tryptone | + | H<sub>2</sub>S production | - |
| Nitrate reduction | + | Gelatin liquefaction | + |
| (+: Positive; -: Negative) |  |  |  |

**Table 5:** Antagonistic activity of *B. amyloliquefaciens* against *S. rubidaea* measured in terms of diameter of the inhibition zone (mean±SD)

| Test organism | Antagonistic activity<br>(Diameter of zone of inhibition in mm) |  |  |  |  |  |  |
| --- | --- | --- | --- | --- | --- | --- | --- |
|  | *B. amyloliquefaciens* (µl/disc) |  |  |  |  |  |  |
|  | 2 | 5 | 10 | 15 | 20 | 25 | 30 |
| *S. rubidaea* | 8.0±0.22 | 9.0±0.42 | 11.0 ±0.23 | 13.0±0.38 | 17.5±0.19 | 24.0±0.44 | 29.0±0.32 |

**Table 6:** Minimum inhibitory concentration (MIC) and minimum bactericidal concentration (MBC) of *B. amyloliquefaciens* against *S. rubidaea*.

| Test organism | Bacterial growth in Muller-Hinton broth<br>(*B. amyloliquefaciens* bacterial concentration) |  |  |  |  |  |  |  |
| --- | --- | --- | --- | --- | --- | --- | --- | --- |
|  | 50 | 100 | 200 | 300 | 400 | 500 | MIC (µl/ml) | MBC (µl/ml) |
| *S. rubidaea* | + | + | - | - | - | - | 200 | 300 |

(+): Growth appears, (-): No growth

### 3.9 Bio-control of *S. rubidaea* in liquid medium

*B. amyloliquefaciens* was co-cultured with approximately 10^5^-10^6^ CFU/ml of *S. rubidaea* in NB medium. The growth rate of *S. rubidaea* appeared to be decreased at 24 h after incubation when compared to the control (Fig. 07).

**Figure. 7:**
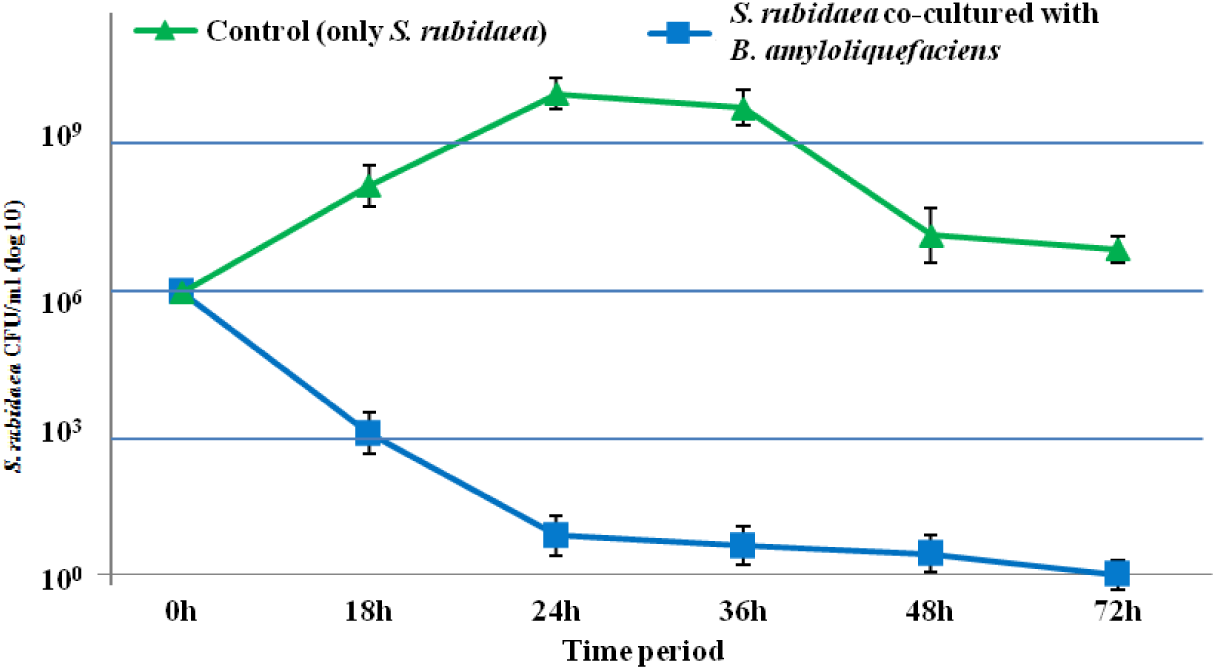
Bio-control by co-culture *S. rubidaea* with *B. amyloliquefaciens*. The colony count of *S. rubidaea* was plotted on the Y axis and time in hours on the X axis.

### 3.10 Antagonistic growth kinetics of *B. amyloliquefaciens* against bacterial pathogens

The previous studied indicating that the strains *B. amyloliquefaciens* possess good antagonistic activity against a broad spectrum of microorganisms such as *Bacillus subtilis, Staphylococcus aureus, Escherichia coli, Pseudomonas aeruginosa* as well as against certain types of fungi [46]. To obtain the antagonistic growth kinetics of *B. amyloliquefaciens* against pathogenic bacteria, nine different human pathogens belonging to both gram-positive and gram-negative were used in this study as test pathogen. The averages diameters of inhibition zones were given in Table 07. The result revealed that *B. amyloliquefaciens* exhibited various degrees of inhibitory activities against the gram-positive and gram-negative pathogenic bacteria, indicating a broad spectrum of inhibition pattern of this organism. Maximum inhibitory activity was observed against *Bacillus cereus* followed by *Bacillus subtilis* and *E. coli*. On the other hand *B. amyloliquefaciens* showed the least susceptibility to *Salmonella entertidis*. Interestingly, previous reports indicating the antibacterial spectrum of *B. amyloliquefaciens* about closely related gram-positive bacteria only, while no inhibitory activity was observed against gram-negative bacteria, especially *E. coli* [23, 47]. Nonetheless, in the present study we found that the *B. amyloliquefaciens* showed antagonistic activity against both gram-positive and gram-negative bacteria. The mode of action (bacteriostatic or bactericidal) of *B. amyloliquefaciens* isolate on test organisms was also determined by inoculating from zone of inhibition on NA. Although *B. amyloliquefaciens* exhibits wide range of zone of inhibition but it was bacteriostatic for most of the pathogen. The above result indicating that the isolate might produce some substances that have inhibitory effect for a wide range of bacteria.

**Table 7:** Antimicrobial activity of *B. amyloliquefaciens* against test pathogens, inhibition zone measured in mm (mean±SD)

| Test organism | Zone of Inhibition (mm) | Mode of Action |
| --- | --- | --- |
| **Gram-positive pathogenic bacteria** |  |  |
| *Bacillus cereus* | 27.0±0.32 | Bc |
| *Bacillus subtilis* | 25.5±0.22 | Bc |
| *Staphylococcus aureus* | 21.5±0.24 | Bc |
| *Enterococcus faecalis* | 22.0±0.30 | Bs |
| **Gram-negative pathogenic bacteria** |  |  |
| *Pseudomonas aeruginosa* | 20.0±0.26 | Bs |
| *Salmonella typhi* | 20.5±0.21 | Bs |
| *Shigella flexinery* | 21.0±0.24 | Bs |
| *Vibrio parahaemolyticus* | 22.5±0.31 | Bs |
| *Escherichia coli* | 24.0±0.28 | Bs |

(Bc= Bactericidal, Bs= Bacteriostatic)

### 3.11 Partial analysis of bacterial metabolites

The secondary metabolites (protein or enzyme or others) of *B. amyloliquefaciens* were determined by SDS-PAGE (10% gel) and was compared with the standard protein marker obtained from Promega, USA (Fig. 11). In 2003, Peng *et al*. reported that *B. amyloliquefaciens* produce a strongly fibrinolytic enzyme called subtilis in DFE which can be used in the treatment of Thrombo-embolism is a complication of medical diseases and surgical procedures [48]. This fibrinolytic enzyme located at **“**31 KDa**”**. In this present study, we also noticed purified enzyme on SDS-PAGE with a molecular weight of approximately 31 KDa which is concurrence with the previous results.

**Figure 11:**
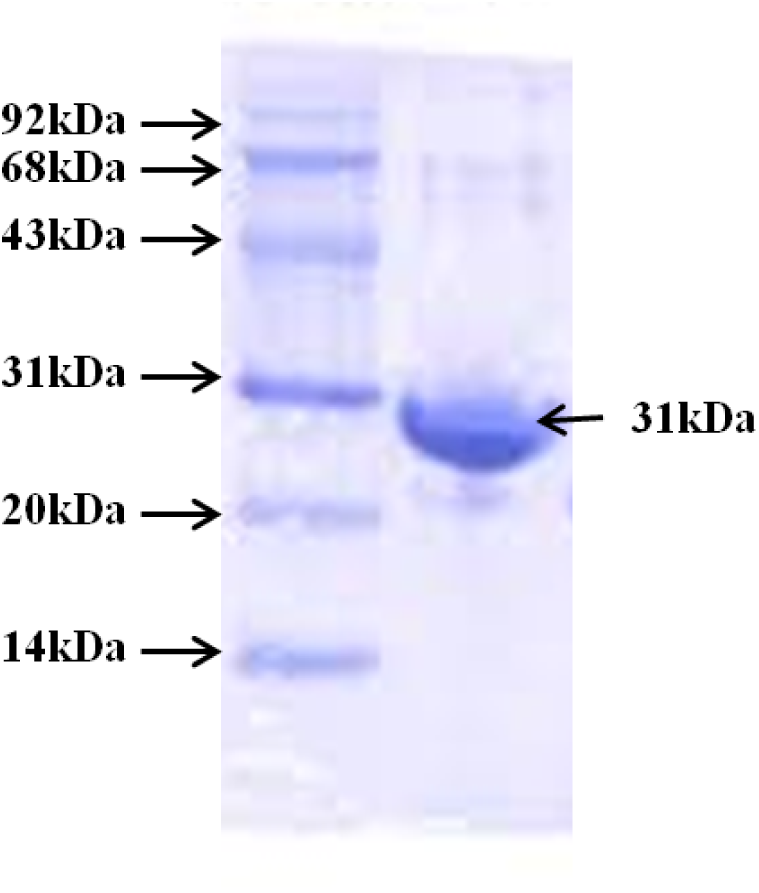
SDS-PAGE analysis.

### 4.0 Conclusion

The numbers of multidrug resistant microbial strains are continuously increasing. This situation provided the impetus to the search for new antimicrobial substances to control multidrug resistant organisms. Several microorganisms have provided a source of hope for novel drug compounds. Results obtained from this study indicated that, unique characteristics of *B. amyloliquefaciens* exhibit strongest antimicrobial activity than the commercially available antibiotics. For instance, Ciproflaxacin showed the maximum zone of inhibition (20 mm) against *S. rubidaea*, where the *B. amyloliquefaciens* showed the maximum zone of inhibition (29 mm) against *S. rubidaea*. Moreover, *B. amyloliquefaciens* can also inhibit a broad range of other pathogenic bacteria. Further genomic analysis about *B. amyloliquefaciens* would be useful for prevention and control *S. rubidaea* as well as will be good candidates for the development of new antimicrobial drugs.

## Data Availability Statement

The experimental data used to support the findings of this study are available from the corresponding author upon request.

## References

1. Ursua PR, Unzaga MJ, Melero P, Iturburu I, Ezpeleta C, Cisterna R: *Serratia rubidaea* as an invasive pathogen. Journal of Clinical Microbiology 1996, 34(1):216–217.

2. Bonnin RA, Girlich D, Imanci D, Dortet L, Naas T: Draft genome sequence of the *Serratia rubidaea* CIP 103234T reference strain, a human opportunistic pathogen. Genome Announc 2015, 3(6):e01340–01315.

3. Saito H, Elting L, Bodey GP, Berkey P: *Serratia* bacteremia: review of 118 cases. Reviews of Infectious Diseases 1989, 11(6):912–920.

4. Menezes EA, Cezafar FC, Andrade MdSdS, Rocha MVAdP, Cunha FA: Frequency of *Serratia* sp in urine infections of intern patients in the Santa Casa de Misericórdia in Fortaleza. Revista da Sociedade Brasileira de Medicina Tropical 2004, 37(1):70–71.

5. Stock I, Burak S, Sherwood KJ, Grüger T, Wiedemann B: Natural antimicrobial susceptibilities of strains of ‘unusual’ *Serratia* species: *S. ficaria, S. fonticola, S. odorifera, S. plymuthica* and S. *rubidaea*. Journal of Antimicrobial Chemotherapy 2003, 51(4):865–885.

6. Hunt M, Mather A, Sánchez BL, Page A, Parkhill J, Keane J, Harris S: ARIBA: rapid antimicrobial resistance genotyping directly from sequencing reads. Microb Genom 3: e000131. In.; 2017.

7. Bush K: Why it is important to continue antibacterial drug discovery. ASM News-American Society for Microbiology 2004, 70(6):282–287.

8. Das B, Neha Nidhi R, Roy P, Muduli A, Swain P, Mishra S, Jayasankar P: Antagonistic activity of cellular components of *Bacillus subtilis* AN11 against bacterial pathogens. International Journal of Current Microbiology and Applied Sciences 2014, 3(5):795–809.

9. Biradar V, Ashwini S, Lachuriye P, Fatima S, Syed A: Isolation of Soil Bacteria for Potential Production of Antibiotics and their Inhibitory Effect on Growth of Pathogens. Int J Curr Microbiol App Sci 2016, 5(8):514–524.

10. Erginkaya Z, Unal E, Kalkan S: Importance of microbial antagonisms about food attribution. Science against microbial pathogens: communicating current research and technological advances 3rd edition Formatex Research Center Spain 2011, 2:1342–1348.

11. Alpas H, Bozoglu F: The combined effect of high hydrostatic pressure, heat and bacteriocins on inactivation of foodborne pathogens in milk and orange juice. World Journal of Microbiology and Biotechnology 2000, 16(4):387–392.

12. Foster JW, Woodruff HB: Antibiotic substances produced by bacteria. Annals of the New York Academy of Sciences 2010, 1213(1):125–136.

13. Rizk M, Rahman T, Metwally H: Screening of antagonistic activity in different *Streptomyces* species against some pathogenic microorganisms. Journal of Biological Science 2007, 7(8):1418–1423.

14. Torres-Cortés G, Millán V, Ramírez-Saad HC, Nisa-Martínez R, Toro N, Martínez-Abarca F: Characterization of novel antibiotic resistance genes identified by functional metagenomics on soil samples. Environmental Microbiology 2011, 13(4):1101–1114.

15. Armstrong E, Yan L, Boyd KG, Wright PC, Burgess JG: The symbiotic role of marine microbes on living surfaces. Hydrobiologia 2001, 461(1-3):37–40.

16. Grossart HP, Schlingloff A, Bernhard M, Simon M, Brinkhoff T: Antagonistic activity of bacteria isolated from organic aggregates of the German Wadden Sea. FEMS Microbiology Ecology 2004, 47(3):387–396.

17. Pathma J, Rahul G, Kamaraj K, Subashri R, Sakthivel N: Secondary metabolite production by bacterial antagonists. Journal of Biological Control 2011, 25(3):165–181.

18. Shelar SS, Warang SS, Mane SP, Sutar RL, Ghosh JS: Characterization of bacteriocin produced by *Bacillus atrophaeus* strain JS-2. International Journal of Biological Chemistry 2012, 6:10–16.

19. Zhang N, Yang D, Kendall JR, Borriss R, Druzhinina IS, Kubicek CP, Shen Q, Zhang R: Comparative genomic analysis of *Bacillus amyloliquefaciens* and *Bacillus subtilis* reveals evolutional traits for adaptation to plant associated habitats. Frontiers in microbiology 2016, 7:2039.

20. Scholz R, Vater J, Budiharjo A, Wang Z, He Y, Dietel K, Schwecke T, Herfort S, Lasch P, Borriss R: Amylocyclicin, a novel circular bacteriocin produced by *Bacillus amyloliquefaciens* FZB42. Journal of Bacteriology 2014, 196(10):1842–1852.

21. Fan B, Blom J, Klenk HP, Borriss R: *Bacillus amyloliquefaciens, Bacillus velezensis*, and *Bacillus siamensis* form an “operational group *B. amyloliquefaciens*” within the *B. subtilis* species complex. Frontiers in microbiology 2017, 8:22.

22. Cao H, He S, Wei R, Diong M, Lu L: *Bacillus amyloliquefaciens* G1: a potential antagonistic bacterium against eel-pathogenic Aeromonas hydrophila. Evidence-Based Complementary and Alternative Medicine, 2011.

23. Boottanun P, Potisap C, Hurdle JG, Sermswan RW: Secondary metabolites from *Bacillus amyloliquefaciens* isolated from soil can kill *Burkholderia pseudomallei*. AMB express 2017, 7(1):16.

24. Borriss R, Chen XH, Rueckert C, Blom J, Becker A, Baumgarth B, Fan B, Pukall R, Schumann P, Spröer C: Relationship of *Bacillus amyloliquefaciens* clades associated with strains DSM 7T and FZB42T: a proposal for *Bacillus amyloliquefaciens* subsp. *amyloliquefaciens* subsp. nov. and *Bacillus amyloliquefaciens* subsp. *plantarum* subsp. nov. based on complete genome sequence comparisons. International journal of systematic and evolutionary microbiology 2011, 61(8):1786–1801.

25. Erkus O: Isolation, Phenotypic, and Genotypic Caharacterization of Yoghurt Starter Bacteria, Master of Science thesis. Food Engineering Graduate School of Engineering and Sciences of Izmir Institute of Technology Izmir, Turkey 2007.

26. Stoyanova M, Bogatzevska N: Phytopathogenic Serratia rubidaea isolated from tulips.

27. Salehi TZ, Mahzounieh M, Saeedzadeh A: Detection of invA gene in isolated *Salmonella* from broilers by PCR method. International Journal of Poulty Science 2005, 4(8):557–559.

28. Charteris WP, Kelly PM, Morelli L, Collins JK: Antibiotic susceptibility of potentially probiotic Lactobacillus species. Journal of Food Protection 1998, 61(12):1636–1643.

29. Lee NK, Yeo IC, Park JW, Kang BS, Hahm YT: Isolation and characterization of a novel analyte from *Bacillus subtilis* SC-8 antagonistic to *Bacillus cereus*. Journal of Bioscience and Bioengineering 2010, 110(3):298–303.

30. Schillinger U, Lücke FK: Antibacterial activity of *Lactobacillus sake* isolated from meat. Applied and Environmental Microbiology 1989, 55(8):1901–1906.

31. Lalpuria M, Karwa V, Anantheswaran RC, Floros J: Modified agar diffusion bioassay for better quantification of N isaplin®. Journal of Applied Microbiology 2013, 114(3):663–671.

32. Jiang L, He L, Fountoulakis M: Comparison of protein precipitation methods for sample preparation prior to proteomic analysis. Journal of Chromatography A 2004, 1023(2):317–320.

33. Lategan M, Torpy F, Gibson L: Control of saprolegniosis in the eel Anguilla australis Richardson, by Aeromonas media strain A199. Aquaculture 2004, 240(1-4):19–27.

34. Hanafy AM, AlMutairi AA, Al-Reedy RM, AlGarni SM: Phylogenetic affiliations of *Bacillus amyloliquefaciens* isolates produced by a bacteriocin-like substance in goat milk. Journal of Taibah University for Science 2016, 10(4):631–641.

35. Gentille D, Perez M, Centelles M: Bacteremia by a *Serratia rubidaea* with an atypical quinolones resistance phenotype. Revista chilena de infectologia: organo oficial de la Sociedad Chilena de Infectologia 2014, 31(3):351–352.

36. Sekhsokh Y, Arsalane L, El MO, Doublali T, Bajjou T, Lahlou IA: *Serratia rubidaea* bacteremia. Medecine et Maladies Infectieuses 2007, 37(5):287–289.

37. Valgas C, Souza SMd, Smânia EF, Smânia Jr A: Screening methods to determine antibacterial activity of natural products. Brazilian Journal of Microbiology 2007, 38(2):369–380.

38. Mannanov R, Sattarova R: Antibiotics produced by *Bacillus* bacteria. Chemistry of Natural Compounds 2001, 37(2):117–123.

39. Kathiresan K, Manivannan S: Amylase production by *Penicillium fellutanum* isolated from mangrove rhizosphere soil. African journal of Biotechnology 2006, 5(10).

40. Kanimozhi M, Johny M, Gayathri N, Subashkumar R: Optimization and production of amylase from halophilic *Bacillus* species isolated from mangrove soil sources. Journal of Applied and Environ Microbiol 2014, 2:70–73.

41. Sen S, Veeranki VD, Mandal B: Effect of physical parameters, carbon and nitrogen sources on the production of alkaline protease from a newly isolated *Bacillus pseudofirmus* SVB1. Annals of Microbiology 2009, 59(3):531–538.

42. Gupta R, Gigras P, Mohapatra H, Goswami VK, Chauhan B: Microbial α-amylases: a biotechnological perspective. Process Biochemistry 2003, 38(11):1599–1616.

43. Li B, Li Q, Xu Z, Zhang N, Shen Q, Zhang R: Responses of beneficial *Bacillus amyloliquefaciens* SQR9 to different soilborne fungal pathogens through the alteration of antifungal compounds production. Frontiers in Microbiology 2014, 5:636.

44. Shaekh M, Mondol A, Islam M, Kabir A, Saleh M, Salah Uddin M, Hoque K, Ekram A: Isolation, characterization and identification of an antagonistic bacterium from *Penaeus monodon*. International Journal of Scientific and Engeneering Research 2013, 4(10):254–261.

45. Ratzke C, Gore J: Modifying and reacting to the environmental pH can drive bacterial interactions. Plos Biology 2018, 16(3):e2004248.

46. Matseliukh E, Safronova L, Varbanets L: *Bacillus amyloliquefaciens* subsp. *plantarum* probiotic strains as protease producers. Biotechnologia Acta 2015, 8(2).

47. Ahmed ST, Islam MM, Mun HS, Sim HJ, Kim YJ, Yang CJ: Effects of Bacillus amyloliquefaciens as a probiotic strain on growth performance, cecal microflora, and fecal noxious gas emissions of broiler chickens. Poultry Science 2014, 93(8):1963–1971.

48. Peng Y, Huang Q, Zhang Rh, Zhang YZ: Purification and characterization of a fibrinolytic enzyme produced by *Bacillus amyloliquefaciens* DC-4 screened from douchi, a traditional Chinese soybean food. Comparative Biochemistry and Physiology part b: Biochemistry and Molecular Biology 2003, 134(1):45–52.

